# Whole organism 3D mapping reveals universal branching topology and biophysical optimization governs vascular and nervous system development

**DOI:** 10.64898/2026.04.10.717729

**Authors:** André Forjaz, Marco Costa, Catarina Oliveira, Paul A. Gensbigler, Lucie Dequiedt, Vasco Queiroga, Saurabh Joshi, William Foster, Matthieu Wyart, Marjorie R. Grafe, Owen J.T. McCarty, Gabriel S. Bever, Jamie O. Lo, Brice Menard, Sean X. Sun, Ashley L. Kiemen, Denis Wirtz

## Abstract

The vascular and nervous systems are transport networks essential for life, yet whether universal geometric and topological principles govern their formation remains unclear. The organism-wide molecular and biophysical coordination required to build and maintain these networks during embryonic development has inspired decades of theoretical work, offering predictions about their expected organization. However, the lack of complete three-dimensional (3D) data has limited validation to isolated structures, leaving whole-organism networks unexplored. Here, we developed a computational pipeline for whole-organism 3D imaging to reconstruct the complete vascular and nervous systems of rhesus macaque, mouse, and turtle embryos. Our analysis reveals that both networks share structural principles, including binary branching and scale-invariant bifurcation geometry, maintained across species and throughout development. Yet, from these shared rules emerge fundamentally different architectures. Vasculature exhibits fractal topology with a fractal dimension ∼3, forming space-filling trees that prioritize proximity to every cell in the body. Nervous system networks exhibit a fractal dimension ∼2, forming sheet-like arbors that prioritize electrical signal transmission. This architectural divergence originates from distinct biophysical constraints operating in bifurcations, where vascular junctions minimize energy expenditure while conserving fluid flow and nerve junctions maximize conduction velocity while conserving electrical current. These local optimization rules, iterated across generations, construct organism-wide networks governed by distinct physical constraints, revealing how evolution generated different solutions for fluid versus electrical transport.

## INTRODUCTION

Angiogenesis and neurogenesis are fundamental morphogenetic processes that establish the vascular and peripheral nervous systems essential for embryonic viability^1–6^. These networks emerge through coordinated cellular behaviors including endothelial tube formation, axonal pathfinding, and branching morphogenesis, creating hierarchical transport architectures to efficiently distribute resources throughout the three-dimensional embryo^7–10^. Understanding the quantitative laws governing vascular and nerve network topology is not only crucial for establishing laws of normal developmental mechanisms, but also for providing a reference framework to study pathologies including congenital vascular malformations, peripheral neuropathies, and tumor angiogenesis^11–14^.

A plethora of theoretical models^15–22^ have speculated on the possible topology (e.g., degree of local branching) and geometry (e.g. local radius) of blood vessels, but a paucity of data due to the lack of comprehensive three-dimensional (3D) reconstructions of whole organisms has greatly limited their validation^23–32^. Consequently, while conformity of vascular and nervous system networks to theoretical predictions has been explored in isolated systems^33–36^, comprehensive cross-species analyses are scarce and existing studies have produced conflicting results, leaving the question of universal conservation unresolved.

Here, we developed a novel automated pipeline to exhaustively segment and skeletonize entire vascular and peripheral nervous systems of whole embryos at high spatial resolution (Fig. 1). We applied this approach to rhesus macaque, mouse, and turtle embryos, enabling comprehensive morphometric analysis of organism-wide transport networks across different amniote lineages at comparable organogenetic stages.

**Fig. 1.**
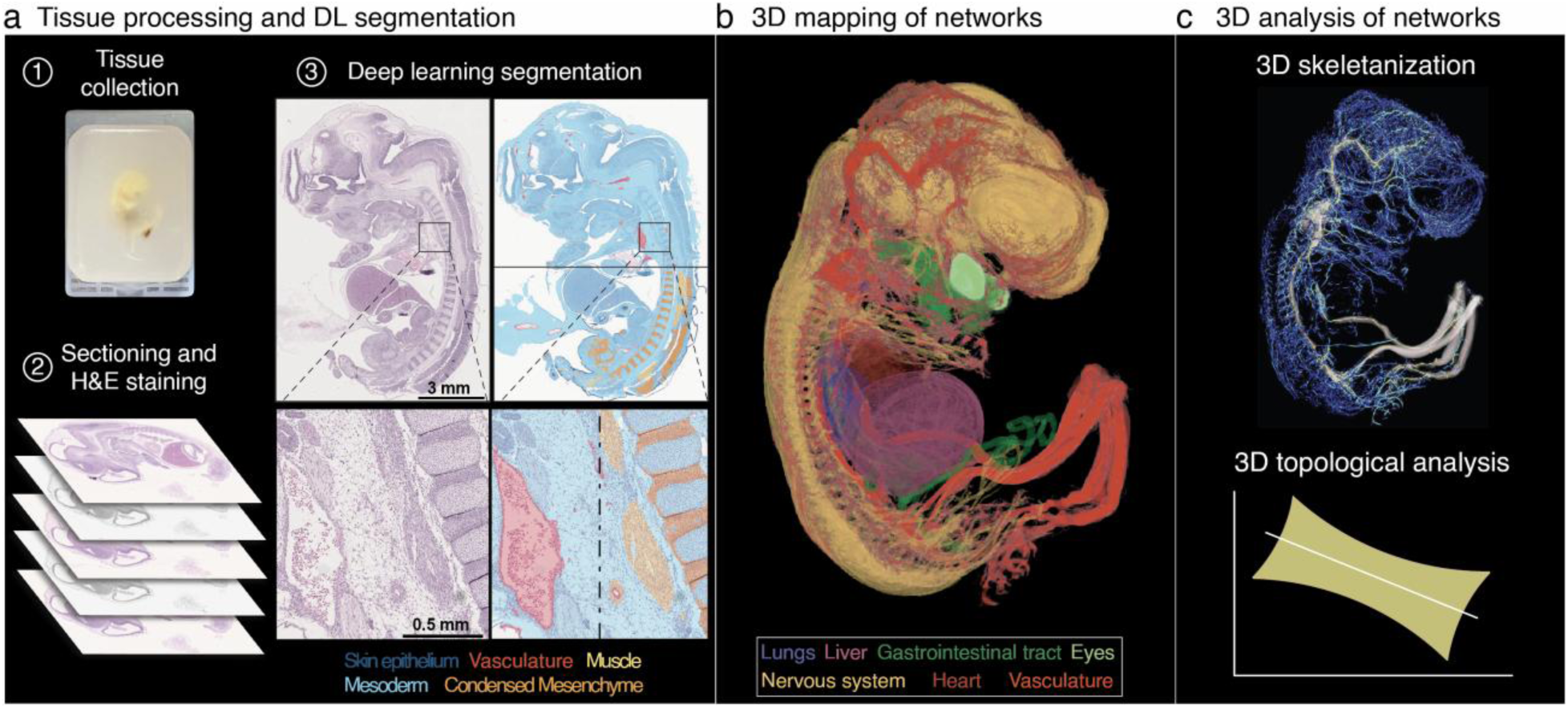
A novel workflow for whole-embryo analysis of vascular and neurvous system networks. **(a)** Embryonic specimens from multiple species were formalin-fixed, paraffin-embedded, and serially sectioned at 4μm thickness. A representative rhesus macaque embryo is shown. Every section was stained with hematoxylin and eosin (H&E) and digitized at 0.5μm/pixel resolution. Deep learning models performed automated pixel-wise semantic segmentation to identify vascular endothelium and lumen, nervous tissue, and extracellular matrix structures. **(b)** Three-dimensional reconstruction and network mapping. Segmented sections were aligned using rigid-elastic registration to generate coherent 3D volumes. Color-coded tissue components include lungs (yellow), liver (orange), gastrointestinal tract (green), eyes (blue), nervous system (black), heart (red), and vasculature (red). **(c)** Graph-theoretical analysis and morphometric extraction. 3D skeletonization using the TEASAR algorithm extracted centerline representations of vascular and nerve networks, enabling quantitative analysis of branching topology, segment lengths, diameter distributions, and hierarchical organization.

Our analysis reveals distinct geometric signatures for each network. Vessel radii follow a power-law with an exponent of -3, while nerve radii scale with an exponent of -2. Remarkably, both systems exhibit log-normal segment length distributions and predominantly binary branching with scale-invariant bifurcation geometry, exhibiting universal characteristics across all examined amniote species. These local branching rules propagate across generations to form fractal architectures over many branching generations. The vasculature exhibits a fractal dimension of ∼3, producing space-filling trees ensuring oxygen delivery to all cells in the embryos. Nerve networks exhibit a fractal dimension of ∼2, forming sheet-like structures that ensure optimized ionic transport for interstitial signaling.

## RESULTS

### Construction of cross-species embryonic datasets for automated vascular and nervous system network analysis

To determine and quantify conserved morphometric principles governing embryonic vascular and nerve development across species, we developed a computational framework integrating serial histopathology with deep learning-based segmentation for whole-organism imaging. We collected embryonic specimens of monkey *Macaca mulatta* (gestational age of 33 and 40 days), mouse *Mus musculus* (gestational age of 11 days), and turtle *Trachemys scripta* (gestational age of 30 and 34 days) (Fig. 2a). These specimens were chosen to provide evolutionarily distinct models for comparison that represent a diversity of developmental timepoints, structural and physiological strategies, and phylogenetic lineages, including both mammalian and non-mammalian vertebrates. These gestational stages were specifically chosen to align with equivalent developmental stages across species, enabling meaningful cross-species morphometric analysis. Specimens were formalin-fixed, paraffin-embedded (FFPE) and serially sectioned at a thickness of four micrometers, with every section stained using hematoxylin and eosin (H&E) and digitized^37^.

**Fig. 2.**
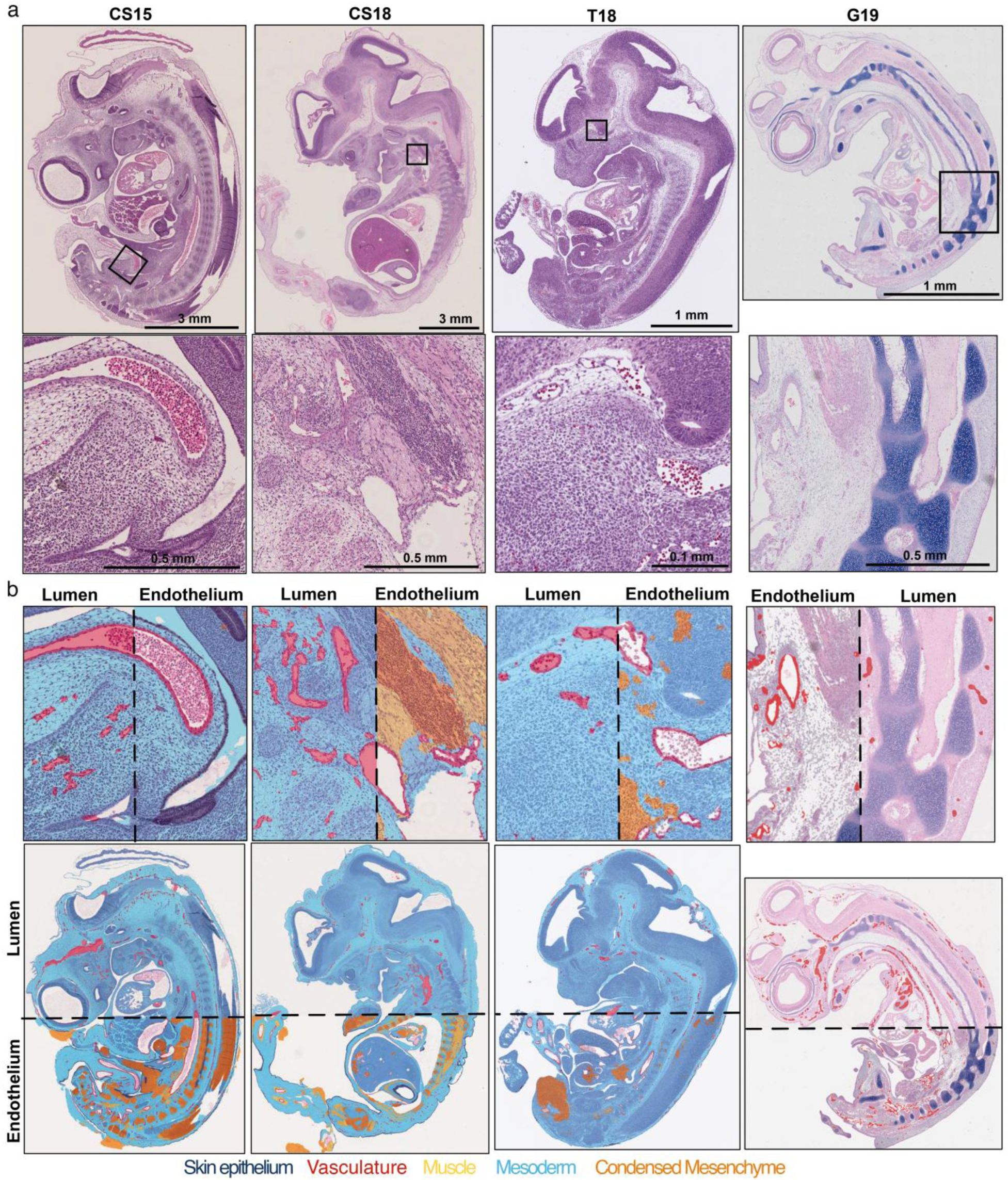
Deep learning segmentation enables automated detection of vascular structures across diverse developmental stages. **(a)** Representative H&E-stained sections from rhesus macaque (CS15 and 18), mouse (T18), and turtle (G19) embryos. Boxed regions highlight areas of detailed segmentation analysis. **(b)** Semantic segmentation of tissue components including lumen (red), endothelium (red walls), muscle (yellow), mesoderm (light blue), and condensed mesenchyme (orange).

Vascular and nervous system structures were labelled using a semantic segmentation algorithm. Vascular training data consisted of 6,250 manually annotated endothelial structures and 6,096 luminal annotations across 122 representative whole slide images (Fig. S1a-d). The segmentation pipeline identified blood vessel endothelium, vessel lumens, from large arterial trunks to capillaries (Fig. 2b). The lumen segmentation model achieved 93.9% accuracy, and the endothelium model reached 96.5% accuracy in an independent test dataset (Fig. S1e,f). We extended this validated segmentation framework to label the developing nervous systems of all specimens. By applying the same training and annotation approach across all H&E-stained serial sections, we successfully labelled nerves, spinal cords, brains and ganglia (Fig. 3).

**Fig. 3.**
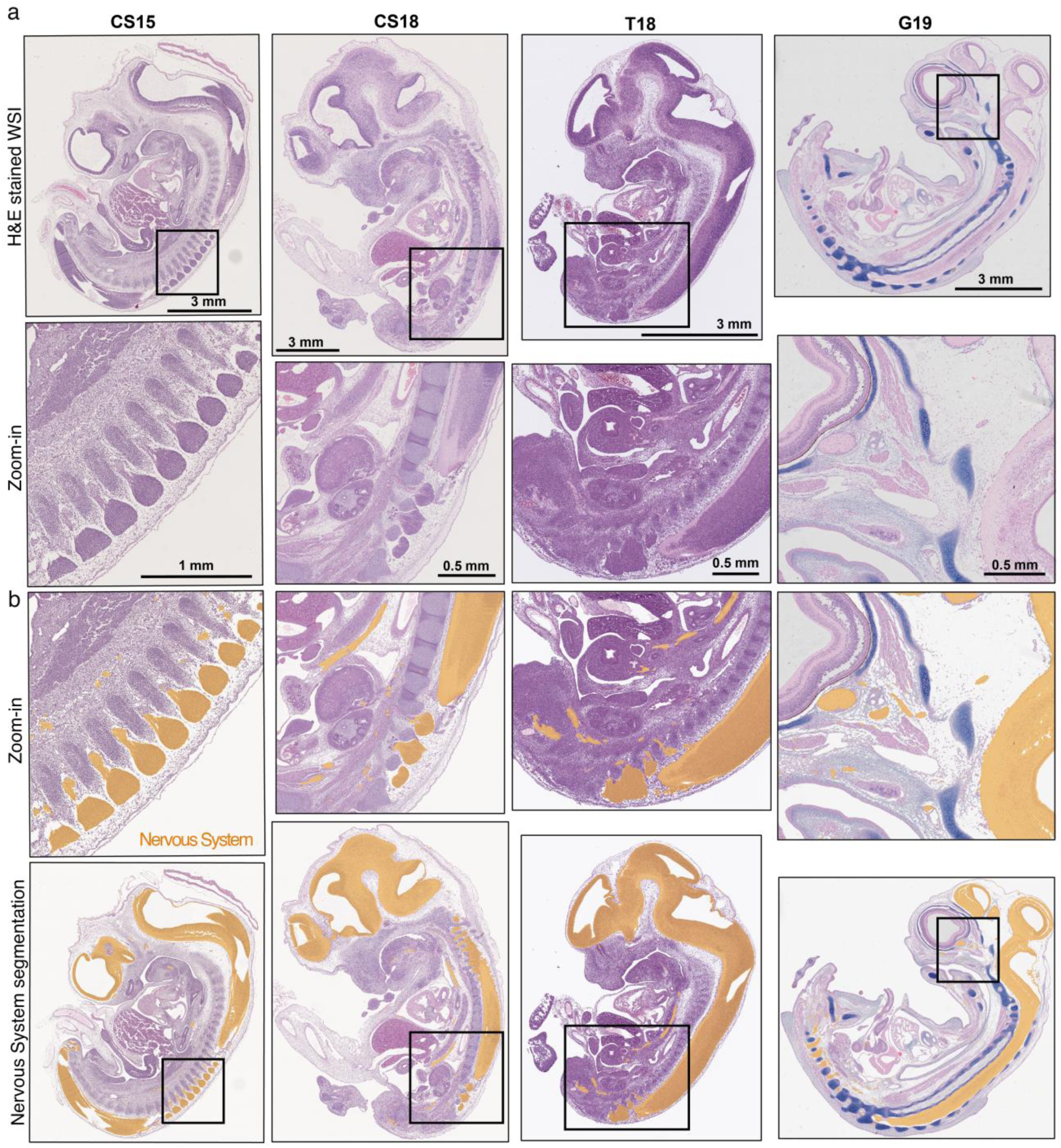
Deep learning segmentation enables automated detection of nervous system microanatomical structures across diverse species and stages. **(a)** Representative H&E-stained sections from rhesus macaque (CS15 and 18), mouse (T18), and turtle (G19) embryos. Boxed regions highlight areas of detailed segmentation analysis. **(b)** Semantic segmentation of nervous system tissue components (orange).

Reconstructing three-dimensional networks from serial histological sections presents significant technical challenges. Registration errors between successive image pairs propagate throughout the stack, causing cumulative distortions that compromise downstream morphometric analysis. To address this, we developed a nonlinear image registration pipeline that aligns H&E images while preserving local tissue architecture, generating continuous volumetric stacks from discrete tissue sections. Registration accuracy was validated using two complementary metrics. First, we calculated pairwise two-dimensional cross-correlation coefficients between successive registered images, requiring values exceeding 0.8 to ensure sufficient structural alignment. Second, we quantified post-registration image warp to detect excessive local deformations, accepting only values below 20% to preserve native tissue geometry (Fig S2). These stringent quality thresholds ensured that registration artifacts did not confound subsequent network analysis.

The combination of exhaustive validated segmentation and high-quality registration yielded accurate digital volumes of the complete vascular and nervous systems across all tested specimens, enabling comprehensive morphometric analysis of organism-wide transport networks.

### Three-dimensional reconstruction enables organism-wide network mapping and reveals hierarchical network organization

Topology refers to the connectivity and branching structure of a network. Geometry refers to the spatial properties of individual network elements, including segment lengths, branch diameters. Our 3D reconstructions constitute the basis to determine the geometry and topology of vascular and nervous systems within developing embryos at single-cell resolution (Fig. 4, Sup. Video 1). Vasculature was 3D-rendered for each specimen to visualize the spatial distribution of major vascular branches and their interconnections (Fig. 4a). This volumetric rendering demonstrated the spatial relationship between the dorsal aorta, cardinal veins, and intersegmental vessels. Vascular networks exhibited hierarchical organization, with large vessels branching progressively into smaller arterioles and capillaries (Fig. 4a). The developing nervous system was similarly 3D-rendered to visualize the neural tube, cranial nerves, and peripheral nerve networks (Fig. 4b). Nerve networks showed distinct patterns of axonal fasciculation and the spatial relationship between central and peripheral nervous system components (Fig. 4b).

**Fig. 4.**
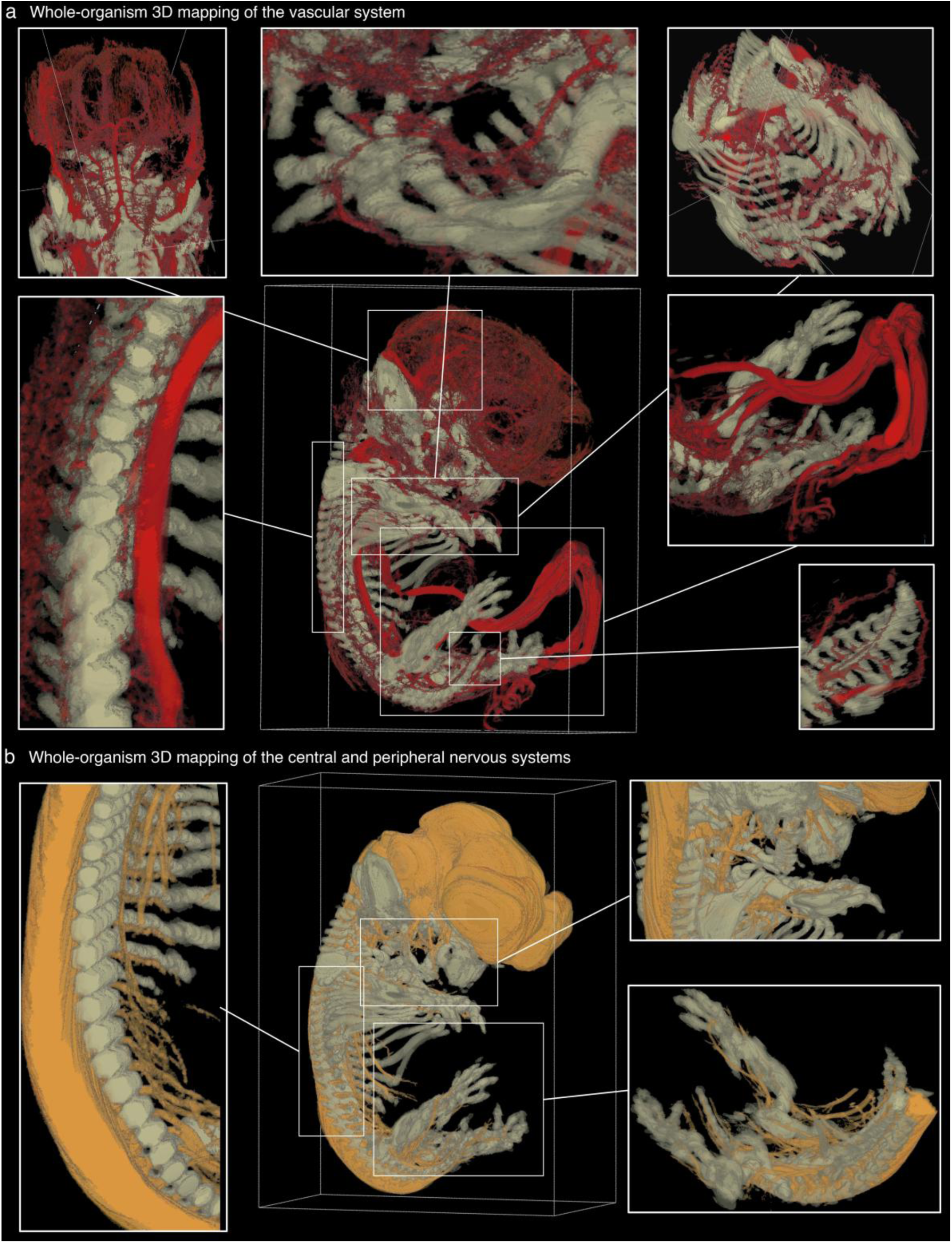
3D mapping of vascular and nervous systems of whole organism at cellular resolution. **(a)** The workflow demonstrates organism-scale, label-free 3D mapping of vasculature and innervation from routine H&E. Rhesus macaque fetus at 40 days gestational age was processed with our deep learning pipeline, showing 3D vascular map (red) and skeletal system (grey). **(b)** 3D segmentation of nerves (orange) shows cranial nerves and spinal cord nerves.

To extract centerline representations of vascular and nerve networks, graph skeletonization was performed, allowing for a quantitative analysis of network topology and geometry (Fig. 5a, 6a). Each network for each embryo was converted to a series of nodes connected by segments, with associated measurements of length, diameter, and spatial coordinates (Table S1). We implemented a TEASAR-based skeletonization approach^38^, which iteratively identifies centerlines by computing the shortest paths through the distance transform of segmented structures. This method maintains centerline accuracy across scales, from large primary vessels to fine capillary networks, while avoiding spurious branches that arise from surface irregularities^39^.

**Fig. 5.**
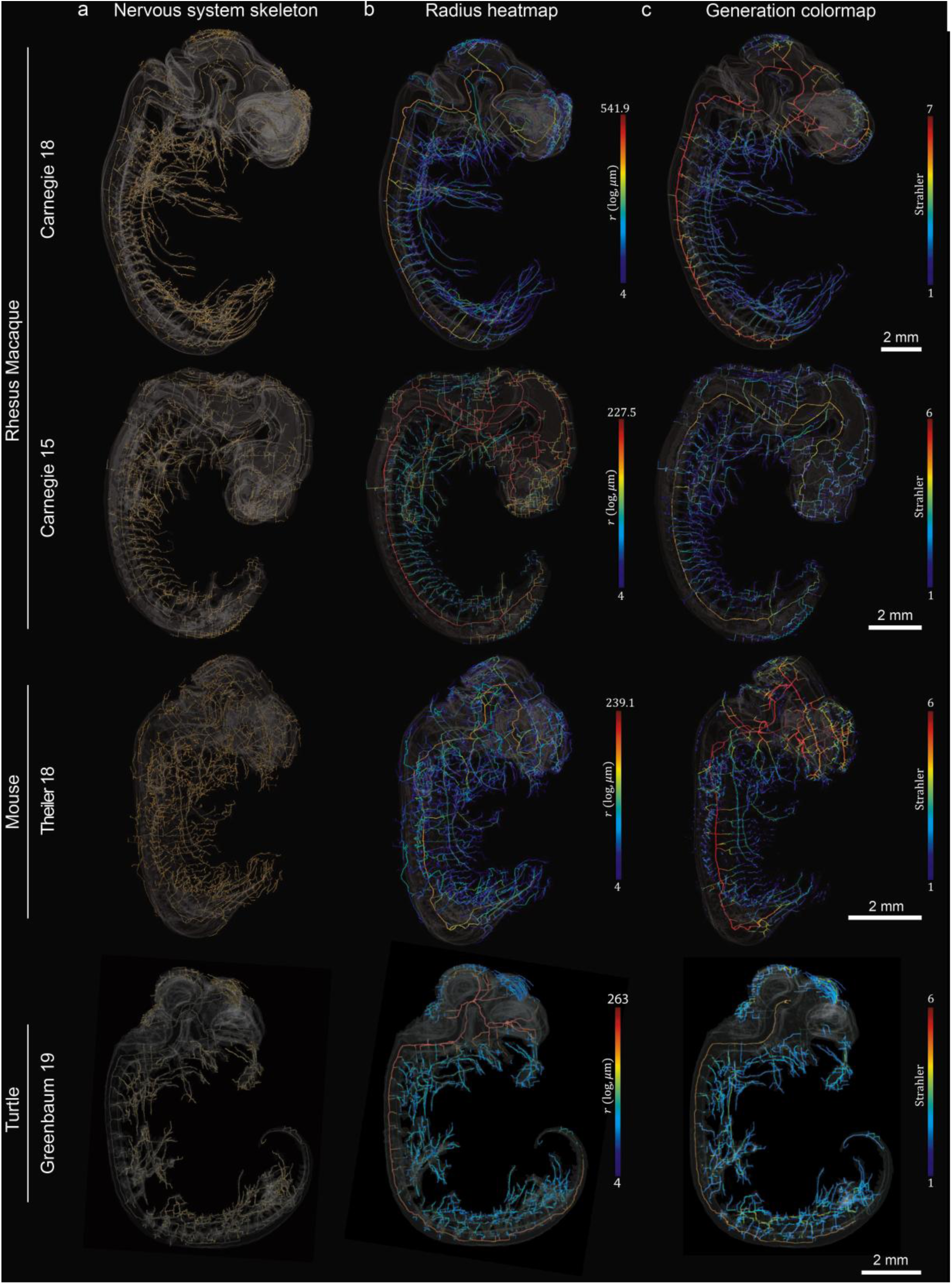
Cross-species three-dimensional reconstruction reveals conserved nervous system network architecture despite phylogenetic diversity. **(a)** 3D skeletonization of nerve networks from rhesus macaque (top), mouse (middle), and turtle (bottom) embryos show hierarchical branching organization across species. **(b)** 3D mapping of radial dimensions for nerve skeletons. **(c)** 3D mapping of branching generations for nerve skeletons.

The calculation of branch radii revealed the hierarchical organization of network diameters, with warm colors indicating larger vessels or nerves and cool colors representing smaller branches (Figs. 5b, 6b). This hierarchical structure reflects the progressive decrease in diameter from proximal to distal segments, where large primary branches subdivide into successively smaller daughter branches. To quantify this hierarchical organization, we applied Strahler ordering, a classification system originally developed for river networks that assigns generation numbers based on branching topology^40^. Terminal branches are assigned order one, and when two branches of equal order merge, the parent branch receives an order incremented by one. This approach provides an objective hierarchical classification independent of absolute size, capturing the nested architecture of biological networks^41^. Strahler order colormaps revealed systematic organization of branching levels across both network types (Figs. 5c, 6c, Sup. Video 2, Sup. Video 3). Higher-generation branches (warm colors) corresponded to major vascular vessels and primary nerves, while lower generations (cool colors) represented peripheral capillaries and terminal axonal branches. The maximum Strahler orders (the number of branching generations) ranged between 7 and 9 for vascular networks, and between 6 and 7 for nerve networks (Fig. 4, 6).

**Fig. 6.**
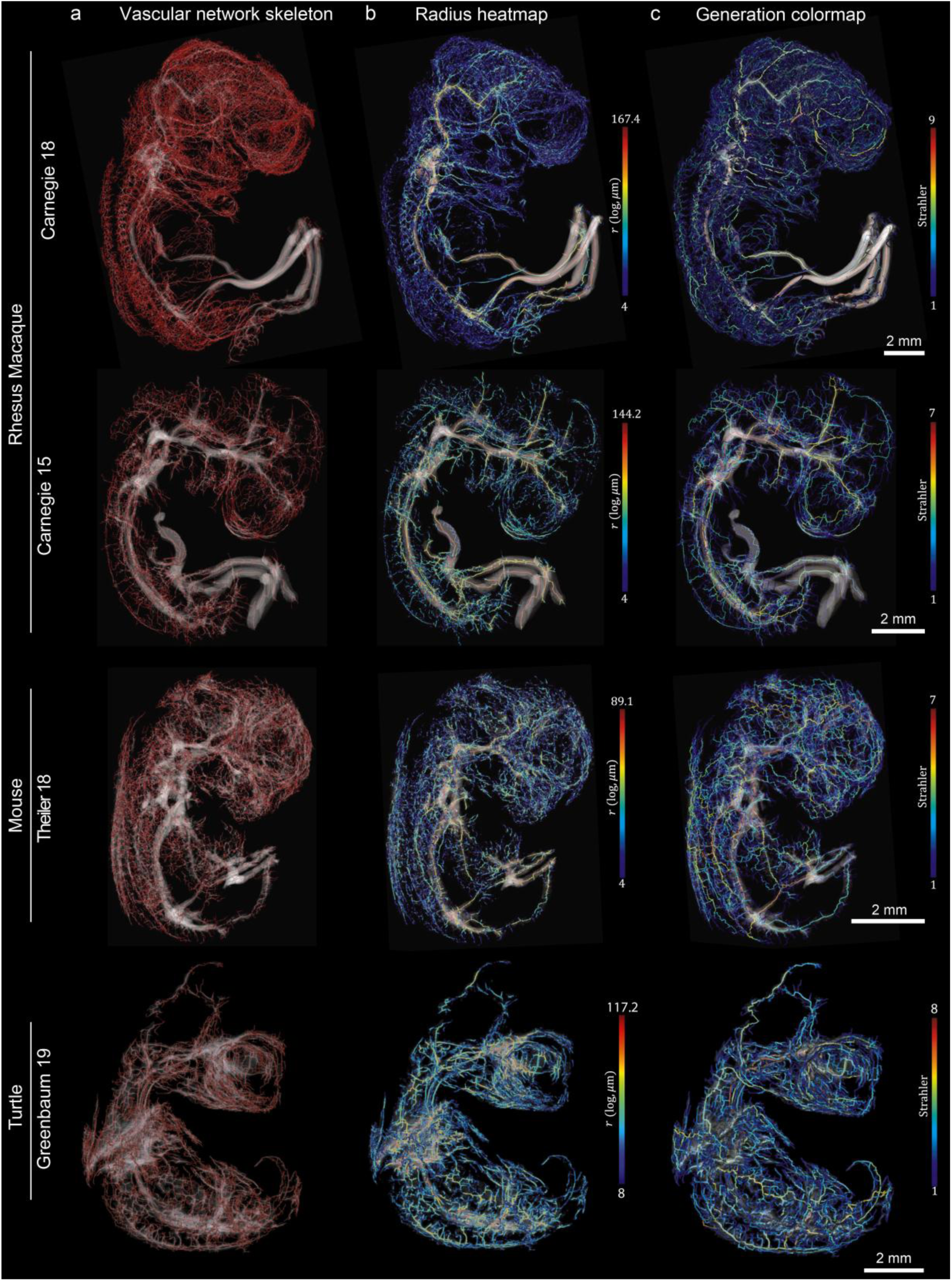
Cross-species three-dimensional reconstruction reveals conserved vascular network architecture despite phylogenetic diversity. **(a)** 3D skeletonization of vascular networks from rhesus macaque (top), mouse (middle), and turtle (bottom) embryos show hierarchical branching organization across species. **(b)** 3D mapping of radial dimensions for vascular skeletons. **(c)** 3D mapping of branching generations for vascular skeletons.

### Vascular networks exhibit conserved scaling principles across species and developmental stages

Three-dimensional analysis of vasculature revealed a conserved scaling law for vessel geometry across species and developmental stages. Indeed, the vessel radius frequency distributions followed power-law scaling (Fig. 7a, Table S2), with exponents derived from CCDF fits (Fig. S4a). Remarkably, these exponents converged to −3 across all specimens: rhesus macaque CS15 (α = −2.99 ± 0.03) and CS18 (α = −3.12 ± 0.02), mouse T18 (α = −3.08 ± 0.04), and turtle G19 (α = −2.60 ± 0.04) and G20 (α = −2.93 ± 0.04). This convergence toward a common exponent of approximately −3 across evolutionary distant species suggests general constraints governing vascular network organization that do not depend on species or developmental stage.

**Fig. 7.**
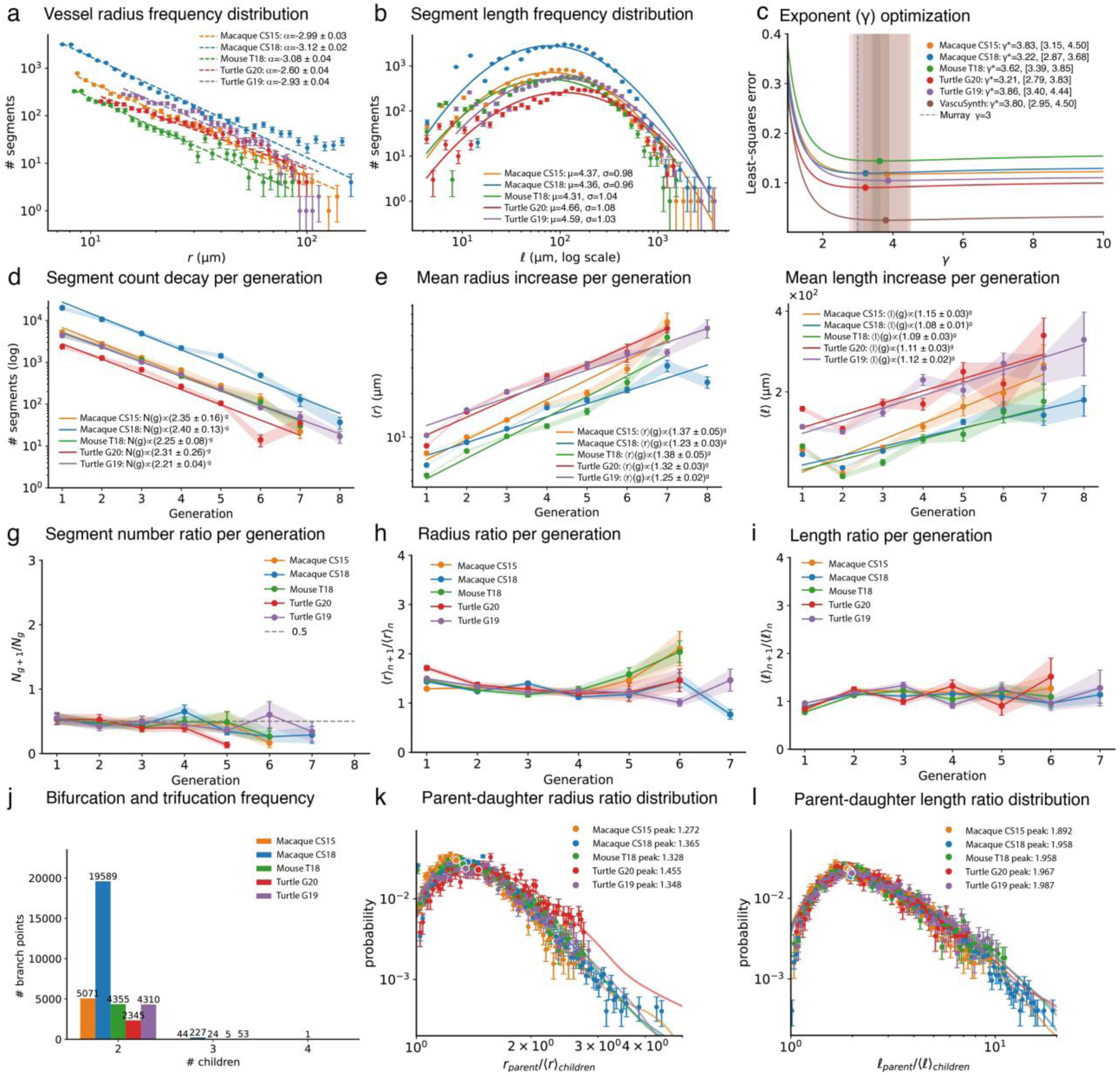
Universal scaling laws govern vascular network organization across species and developmental stages. **(a)** Vessel radius frequency distributions follow POWER statistics with consistent parameters across macaque (CS15, CS18), mouse (T18), and turtle (G19) specimens. **(b)** Segment length distributions exhibit similar log-normal scaling conserved across species. **(c)** Murray’s law exponent (γ) optimization via least-squares error analysis identifies optimal values for each specimen. **(d)** Segment count decays exponentially with branching generation, with consistent scaling exponents across species. **(e)** Mean vessel radius increases systematically per generation, following exponential relationships. **(f)** Radius ratio scaling demonstrates conserved exponential growth across generations. **(g)** Children-to-parent segment number ratios converge toward 0.5, indicating symmetric bifurcation dominance. **(h)** Children-to-parent radius ratios per specimen. **(i)** Length ratios reveal generation-dependent vessel geometry. **(j)** Branch point distribution confirms bifurcation dominance over higher-order branching. **(k)** Parent-to-daughter radius ratio probability distributions cluster near Murray’s law predictions. **(l)** Parent-to-children length ratio distributions demonstrate consistent branching geometry across species. Horton ratios remain constant across generations, indicating scale-invariant hierarchical organization conserved from reptiles to primates.

The distributions of vessel segment lengths were well described by log-normal functions (Fig. 7b, S4b), with parameters that were conserved across species for rhesus macaque CS15 (μ = 4.37, σ = 0.98) and CS18 (μ = 4.36, σ = 0.96), mouse T18 (μ = 4.31, σ = 1.04), and turtle G19 (μ = 4.59, σ = 1.03) and G20 (μ = 4.66, σ = 1.08). This log-normal distribution of segment lengths suggests stochastic growth processes constrained by local tissue architecture, where multiplicative factors during development generate characteristic right-skewed distributions^42^.

### Vascular local branching geometry and biophysical optimization

To understand the underlying geometry of the vasculature across species, we examined individual branching bifurcations by quantifying parent-to-daughter ratios of vessel radii and segment lengths (Fig. 7k, 7l, S4d).

To understand the relationship between parent and daughter vessel radii at bifurcations points, we investigated the exponent γ such that the parent radius raised to γ equals the sum of daughter radii raised to the same power. Least-squared optimization of the branching exponent (γ) confirmed values close to the theoretical optimum of 3 for fluid transport, with rhesus macaque CS15 (γ* = 3.83) and CS18 (γ* = 3.22), mouse T18 (γ* = 3.62), and turtle G19 (γ* = 3.86) and G20 (γ* = 3.21) (Fig. 7c). Synthetic vascular networks generated using VascuSynth^43^, which explicitly implements Murray’s law branching rules, exhibited comparable deviation from the theoretical optimum (γ* = 3.80), confirming that our measured values are consistent with Murray’s law optimization.

Parent-daughter length ratio distribution peaked at ∼2 (Fig. 7l, S4d right), indicating that parent segments were approximately twice the length of daughter segments on average. The mean tortuosity, defined as the ratio of actual path length to Euclidean distance, remained constant at approximately 1.2 across branch generations and species (Fig. S4e), indicating that blood vessels maintain consistent geometry throughout the branching hierarchy. The distribution of parent-daughter radius ratios peaked at 1.26 across all species (Fig. 7k, S4d left).

The quantification of branching topology confirmed that bifurcations dominated across all species, accounting for more than 99% of branch points, while trifurcations and higher-order branching events were rare (Fig. 7j). This predominantly binary branching establishes the foundation for hierarchical organization of vasculature networks. Notably, while segment number ratio histograms peak at 2 reflecting binary physical junctions, Horton’s bifurcation ratio (R_N)_ from segment count decay plots ranges between 2.3-2.4. This discrepancy arises because Strahler ordering allows a parent segment to maintain its generation when joined by lower-order branches, enabling a single higher-order segment to serve as hierarchical parent to multiple side-branches along its length, thereby inflating lower-order segment counts and yielding R_N_ values exceeding 2.

### Local branching rules of the vasculature propagate across generations to form fractal architectures

Having established that individual bifurcations follow conserved optimization principles, we next investigated whether these local rules propagated uniformly across the network hierarchy to generate fractal architectures (Fig. 7d–f). We applied Strahler ordering^40,41^, with peripheral capillaries receiving low generation numbers and central vessels receiving progressively higher values.

Segment count exhibited exponential decay with increasing generation number (Fig. 7d), a hallmark of fractal branching networks^15,44^. The mean branching ratio, calculated from exponential regression of segment count versus generation, converged to ∼2 across all species and developmental stages: rhesus macaque CS15 (n = 2.35 ± 0.16) and CS18 (n = 2.40 ± 0.13), mouse T18 (n = 2.25 ± 0.08), and turtle G19 (n = 2.21 ± 0.04) and G20 (n = 2.31 ± 0.26). This branching ratio quantifies the hierarchical reduction in segment number across generations, with a value of 2 reflecting a predominantly binary branching (Fig. 7j, Table S4).

The mean vessel radius scaled exponentially with the generation number, increasing systematically from peripheral capillaries to central vessels (Fig. 7e). The radius scaling factor at each generation was conserved across species and developmental stages: rhesus macaque CS15 (1.37 ± 0.05) and CS18 (1.23 ± 0.03), mouse T18 (1.38 ± 0.05), and turtle G19 (1.25 ± 0.02) and G20 (1.32 ± 0.03). The mean segment length exhibited a similar exponential scaling: rhesus macaque CS15 (1.15 ± 0.03) and CS18 (1.08 ± 0.01), mouse T18 (1.09 ± 0.03), and turtle G19 (1.12 ± 0.02) and G20 (1.11 ± 0.03) (Fig. 7f, Table S4).

To assess the potential self-similarity nature of the vasculature, a defining property of fractal networks, we computed successive generation ratios for each morphometric parameter (Fig. 7g–i). The segment number ratios showed a constant value of approximately 0.5 across generations (Fig. 7g). Remarkably, both the radius (Fig. 7h) and length ratios (Fig. 7i) between branches of successive generation numbers were similarly generation-invariant, maintaining constant values regardless of hierarchical position. This scale-invariance provides quantitative evidence for a fractal organization of the vasculature, with local bifurcation rules propagating across hierarchical levels to generate fractal networks of fractal dimension 3.

A fractal dimension of 3 indicates that blood vessels are “space-filling”^15,45^, which would minimize vascular-parenchymal diffusion distance, optimizing metabolic exchange (see mathematical derivation in the Supplemental Material). The presence of the same fractal architecture in macaque, mouse, and turtle embryos indicates a deeply conserved scaling relationship that likely reflects a fundamental biophysical optimization

### Nervous system networks exhibit conserved scaling principles across species and developmental stages

Next, we analyzed the three-dimensional geometry and topology of the nervous systems. The distributions of nerve radius frequencies followed power-law scaling, with exponents derived from CCDF fits and converging to −2 across all specimens. Exponents were computed (Fig. 8a, S5a, Table S3) for rhesus macaque CS15 (α = −1.92 ± 0.02) and CS18 (α = −2.09 ± 0.03), mouse T18 (α = −1.87 ± 0.02), and turtle G19 (α = −2.41 ± 0.03) and G20 (α = −2.55 ± 0.07). These exponents are notably shallower than vascular networks (α ≈ −3), indicating a flatter diameter distribution with relatively greater representation of large-caliber nerve fibers.

**Fig. 8.**
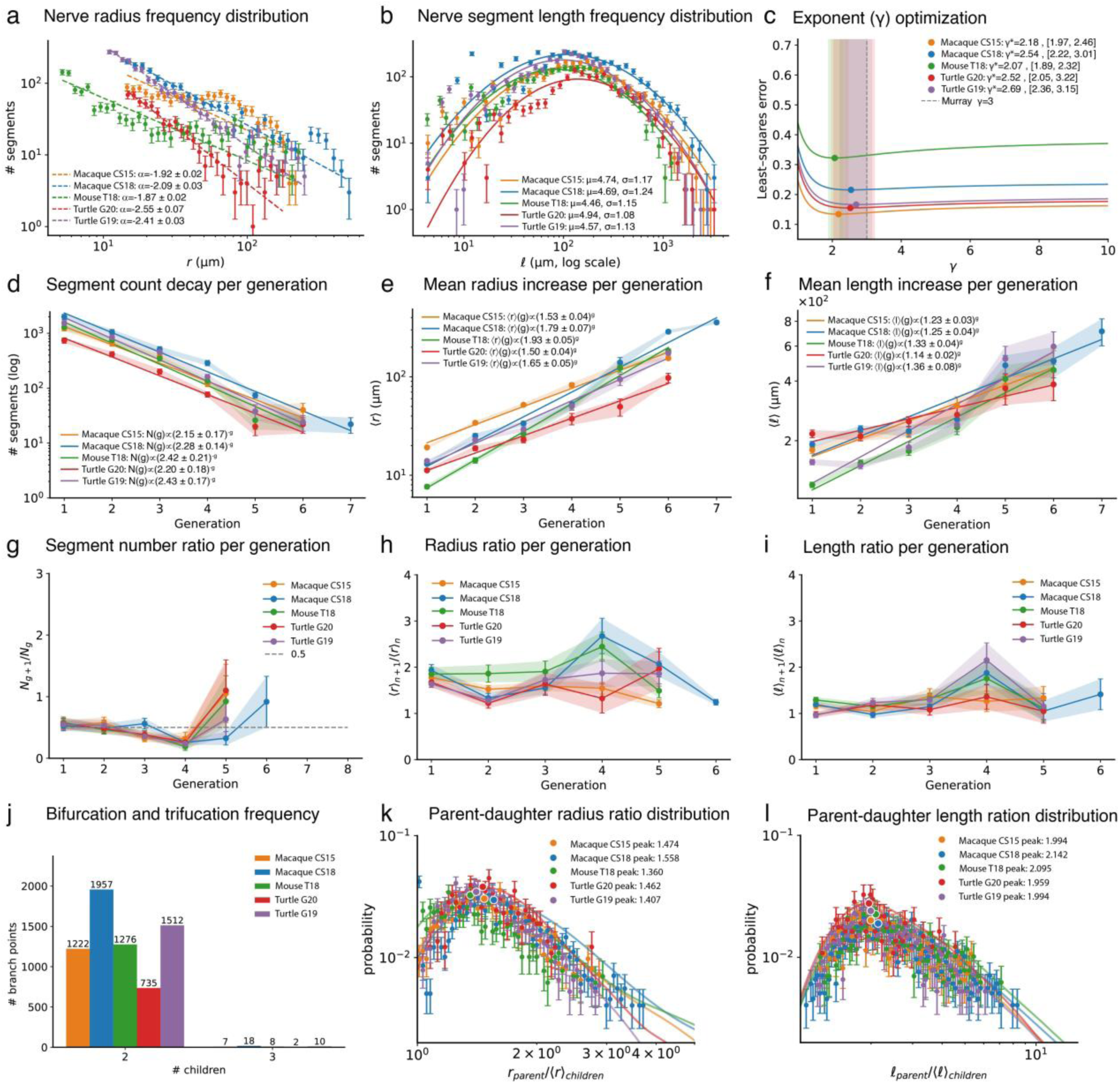
Nervous system networks exhibit distinct scaling properties optimized for electrical conduction. **(a)** Nerve radius frequency distributions follow power-law statistics with consistent parameters across rhesus macaque (CS15, CS18), mouse (T18), and turtle (G20, G19) specimens. **(b)** Segment length distributions exhibit similar log-normal scaling conserved across species. **(c)** Murray’s law exponent (γ) optimization via least-squares error analysis identifies optimal values for each specimen, with nerve networks showing γ values closer to 2.0 compared to the vascular optimum of 3.0. **(d)** Segment count decays exponentially with branching generation, with consistent scaling exponents across species. **(e)** Mean nerve radius increases systematically per generation, following exponential trend. **(f)** Mean nerve length increases per generation, following exponential trend. **(g)** Daughter-to-parent segment number ratios converge toward 0.5, indicating symmetric bifurcation dominance. **(h)** Bifurcation radius ratios demonstrate generation-dependent relationships across nerve hierarchies. **(i)** Length-to-radius ratios reveal generation-dependent nerve geometry. **(j)** Branch point distribution confirms bifurcation dominance over higher-order branching. **(k)** Parent-to-daughter radius ratio probability distributions cluster near Murray’s law predictions. **(l)** Parent-to-daughter length ratio distributions demonstrate consistent branching geometry across species.

Nerve segment length distributions were well described by log-normal functions (Fig. 8b, S5b), with conserved parameters for rhesus macaque CS15 (μ = 4.74, σ = 1.17) and CS18 (μ = 4.69, σ = 1.24), mouse T18 (μ = 4.46, σ = 1.15), and turtle G19 (μ = 4.57, σ = 1.13) and G20 (μ = 4.94, σ = 1.08). Similarly to the vasculature, log-normal segment lengths indicate stochastic growth constrained by local tissue architecture. The higher variance (σ ≈ 1.1–1.2) compared to vascular segments reflects greater heterogeneity in nerve fiber lengths.

### Nervous system bifurcations optimize for electrical signal transmission

To determine the optimization principles underlying these distributions, we analyzed the geometry of nerves at the branching points of the network by quantifying parent-to-daughter radii and length ratios. The parent-daughter radius ratio distribution (Fig. 8k, 8l, S5d), with species-specific peaks for rhesus macaque CS15 (1.474) and CS18 (1.558), mouse T18 (1.360), and turtle G19 (1.407) and G20 (1.462). Least-squared optimization of the branching exponent γ confirmed values near 2, substantially below the vascular optimum of 3: rhesus macaque CS15 (γ* = 2.18) and CS18 (γ* = 2.54), mouse T18 (γ* = 2.07), and turtle G19 (γ* = 2.69) and G20 (γ* = 2.52) (Fig. 8c). Linear regression of parent-daughter radius relationships confirmed a deviation from vascular cubic scaling, with slopes below unity (β = 0.82–0.88, R² = 0.74–0.81; Fig. S5c). A branching exponent near 2 indicates optimization for electrical signal transmission, where bifurcations conserve current while maximizing conduction velocity (see mathematical derivation in the Supplemental Material).

Parent-daughter length ratio distribution peaked near 2 across all species, with values for rhesus macaque CS15 (1.994) and CS18 (2.142), mouse T18 (2.095), and turtle G19 (1.994) and G20 (1.959) (Fig. 8l, S5d right). The mean tortuosity of nerves remained constant at approximately 1.1–1.2 across all generations and species (Fig. S5e), indicating relatively straight fiber trajectories throughout the branching hierarchy. While trifurcations and higher-order branching events were more prevalent than in vascular networks, bifurcations dominated across all species, accounting for the majority of branch points (Fig. 8j). Similar to the vasculature, segment number ratio histograms peak at 2 reflecting binary physical junctions, while Horton’s bifurcation ratio (R_N_) from segment count decay plots ranges between 2.3-2.4, as Strahler ordering allows higher-order segments to serve as hierarchical parents to multiple lower-order side-branches along their length.

The convergence of these distributions suggests that nerve branching geometry follows conserved principles across species. They were distinct from those found for the vasculature, which would reflect the fundamentally different physics of electrical versus fluid transport. While vascular networks optimize for minimizing hydrodynamic resistance (γ ≈ 3), nerve networks appear to be optimized for electrical signal transmission with scaling exponents near γ ≈ 2, consistent with cable theory predictions for maximizing conduction velocity while minimizing material cost^46,47^.

### Local branching rules propagate across generations to form surface-optimized nervous system networks

We next investigated whether these local bifurcation rules propagated uniformly across the network hierarchy (Fig. 8d-f). The segment count decayed exponentially with the generation number (Fig. 8d, Table S5), yielding similar branching ratios across species: rhesus macaque CS15 (n = 2.15 ± 0.17) and CS18 (n = 2.28 ± 0.14), mouse T18 (n = 2.42 ± 0.21), and turtle G19 (n = 2.43 ± 0.17) and G20 (n = 2.20 ± 0.18). Again, these values suggested that binary bifurcation is the dominant branching principle for nervous networks.

The mean nerve radius increased exponentially from peripheral to central nervous system (Fig. 8e), with scaling factors for rhesus macaque CS15 (1.53 ± 0.04) and CS18 (1.79 ± 0.07), mouse T18 (1.93 ± 0.05), and turtle G19 (1.65 ± 0.05) and G20 (1.90 ± 0.04). These radius scaling factors (1.5-1.9) substantially exceed vascular values (1.2-1.4), indicating a steeper radius tapering across nerve generations. The mean segment length increased with generation, with scaling factors for rhesus macaque CS15 (1.23 ± 0.03) and CS18 (1.25 ± 0.04), mouse T18 (1.33 ± 0.04), and turtle G19 (1.36 ± 0.08) and G20 (1.14 ± 0.02) (Fig. 8f, Table S5). These length scaling factors also exceed vascular values, further suggesting that nerve networks employ steeper hierarchical tapering.

Successive generation ratios revealed differences from vascular self-similarity. The segment number ratios trended toward 0.5 across generations (Fig. 8g), consistent with binary branching. However, radius ratios (Fig. 8h) and length ratios (Fig. 8i) exhibited greater generation-dependence than observed for vascular networks, suggesting deviation from strict scale-invariance. Despite reduced self-similarity, conserved local optimization at nerve bifurcations generated networks with fractal dimension approaching 2. This dimension corresponds to a sheet-like topology that is optimized for tissue innervation rather than volumetric space-filling.

The convergence of morphometric properties across macaque, mouse, and turtle embryos demonstrates that vertebrate nervous and vascular networks reflect distinct biophysical optimization. While vascular networks present space-filling topology with fractal dimension of ∼3 (Fig. 6), nervous networks feature sheet-like arborizations with fractal dimension of ∼2 (Fig. 5).

## DISCUSSION

We present a 3D computational framework that integrates serial histopathology, deep learning-based segmentation, and volumetric reconstruction for whole-organism quantitative analysis of embryonic vascular and nervous system networks. This workflow enabled extraction of topological and geometric parameters from thousands of individual network segments. To the best of our knowledge, this study presents the first organism-wide mapping of both vascular and nerve networks at cellular resolution, capable of simultaneously quantifying branching topology, diameter distributions, and scaling relationships across multiple phylogenetic lineages.

The 3D arrangement of embryonic transport networks is subject to several constraints. Given the diversity of processes that govern the development of these networks, we would expect that different functional demands yield different architectural solutions. We show that vascular and nerve networks, despite serving fundamentally different transport functions, share common organizational principles while displaying systematic differences that reflect their specific biophysical requirements. Our results across phylogenetic lineages indicate both vascular and nerve network types exhibit scale-invariant bifurcation ratios of approximately 0.5, with binary branching accounting for more than 99% of vasculature and the majority of nerve branch points. This validates a prediction of WBE metabolic scaling theory^28^ and suggests that symmetric bifurcation represents an evolutionarily conserved solution to hierarchical network construction^49,50^.

Despite these shared organizational principles, vascular and nerve networks diverge in their optimization targets, reflecting the fundamentally different physics of fluid and electrical transport (see Supplemental Material). Vascular networks exhibited radius power-law exponents converging to −3 across all species, revealing a universal scaling law governing vessel diameter distribution. To determine whether this distribution reflected network optimization principles, we examined bifurcation geometry. The branching exponent (γ* = 3.21-3.86) closely approximated the theoretical optimum of γ = 3 predicted for bifurcations that minimize the combined metabolic cost of pumping power and vessel maintenance while conserving volumetric flow^48^. Notably, synthetic vascular networks generated using VascuSynth^43^, which explicitly implements Murray’s law, also exhibited comparable deviation from the theoretical optimum (γ* = 3.80), suggesting that observed deviations reflect measurement variability and finite network effects rather than departures from the underlying optimization principle. Parent-daughter radius ratios peaked close to Murray’s predicted 2^1/3^. These findings provide empirical support for the hypothesis that vascular network evolution minimized metabolic costs subject to flow efficiency constraints.

In contrast, the nervous system networks exhibited radius power-law exponents converging to ∼2 across all species, revealing a universal scaling law for nerve diameter distribution distinct from the vasculature. Examination of bifurcation geometry revealed branching exponents (γ* = 2.07-2.69) clustering near the theoretical optimum of γ = 2 predicted for networks optimizing electrical signal transmission. This exponent aligns with cable theory predictions for unmyelinated axons^46,47^, where conduction velocity scales as the square root of the axon diameter. Importantly, myelination represents a late-developmental event across vertebrates, occurring postnatally in mice, during late second to third trimester in macaques, and near hatching in turtles. At the embryonic stages examined, axons remain entirely unmyelinated, thus validating application of the uniform membrane assumption underlying standard cable theory.

This divergence in local bifurcation optimization propagates across branching generations to produce distinct global network topologies. Vascular networks demonstrated strong self-similarity, with generation-invariant scaling factors for radius (1.23-1.38) and length (1.08-1.15). This scale-invariance across hierarchical levels, combined with branching ratios of 2, generates fractal networks with dimension of 3. Such a resulting space-filling topology minimizes diffusion distance between vasculature and parenchyma, ensuring efficient oxygen delivery to every cell. Nerve networks exhibited steeper radius scaling factors (1.53-1.93) and greater generation-dependence, indicating deviation from strict self-similarity. This reduced scale-invariance, combined with branching exponents of 2, generates networks with fractal dimension approaching 2, producing planar distributions optimized for tissue innervation rather than volumetric coverage. Log-normal segment length distributions in both systems indicate stochastic growth processes constrained by local tissue architecture, where multiplicative developmental factors generate characteristic right-skewed distributions independent of the transported medium.

Recent work showed that surface minimization governs local branching geometry in physical networks^49^. While that analysis focused on local features (branching angles, trifurcation emergence), our generation-resolved approach reveals complementary global hierarchical scaling, suggesting that biological transport networks obey distinct optimization principles across spatial scales. Local bifurcation geometry minimizes surface area and metabolic cost at individual branch points, while these rules iterate across generations to produce organism-wide network organization governed by transport-specific physical constraints.

More empirical studies are needed to validate these scaling predictions across developmental stages. Future work could also expand temporal scope and include molecular characterization of vascular and nerve development to better understand the biochemical mechanisms underlying global and local vascular and nervous developmental processes^50–52^. Additionally, these complete 3D network reconstructions could serve as anatomically accurate templates for bioprinting functional vascular and neural scaffolds, potentially advancing tissue engineering and regenerative medicine applications.

This study demonstrates a first-in-kind scalable platform for embryonic transport network analysis. Universal power-law scaling governs radius distributions in both systems, with distinct exponents reflecting different optimization targets. Vascular bifurcations exhibit branching exponents near three, consistent with minimizing metabolic cost for fluid transport, while nerve bifurcations exhibit exponents near two, consistent with maximizing conduction velocity for electrical transmission. These local optimization rules propagate across branching generations to generate space-filling vascular topology and planar nerves distributions. These principles establish a quantitative framework for understanding normal development and vascular and neural pathologies.

## SUPPLEMENTAL FIGURES

**Fig. S1.**
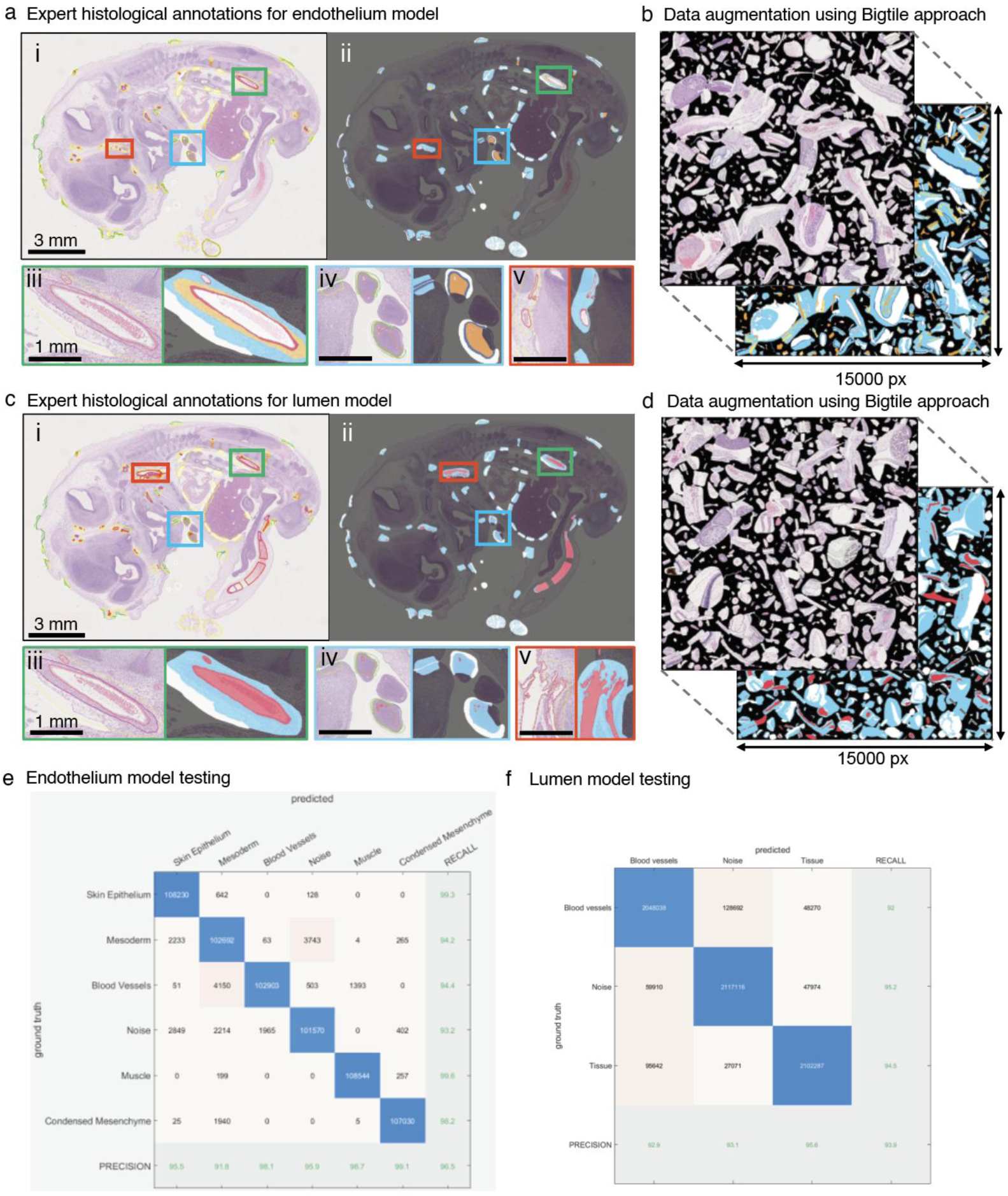
Deep learning model development and validation for automated vascular network segmentation. **(a)** Pathologist expert histological annotations for endothelium model training. (i-ii) Representative H&E-stained embryonic section with color-coded annotation boxes highlighting diverse vascular structures across anatomical regions. (iii-v) Expert-annotated ground truth masks showing pixel-wise endothelium classification (red) overlaid tissue sections. High-magnification detail views of annotated regions demonstrating endothelium identification in various morphological contexts including large vessels, capillary networks, and developing vascular buds. **(b)** Data augmentation using big tile approach, with systematic tiling, to generate training patches while preserving spatial context. **(c)** Pathologist expert histological annotations for lumen model training. (i-ii) Identical embryonic section with annotation boxes highlighting vessel lumens and cavities. (iii-v) Expert-annotated ground truth masks showing lumen classification (red) with clear delineation of vascular spaces. Detail views demonstrating lumen annotation across vessel sizes from major arterial spaces to smaller capillary lumens. **(d)** Big tile data augmentation strategy for lumen model development. **(e)** Endothelium model testing performance demonstrated 96.5% accuracy. **(f)** Lumen model testing performance demonstrated 93.9% accuracy.

**Fig. S2.**
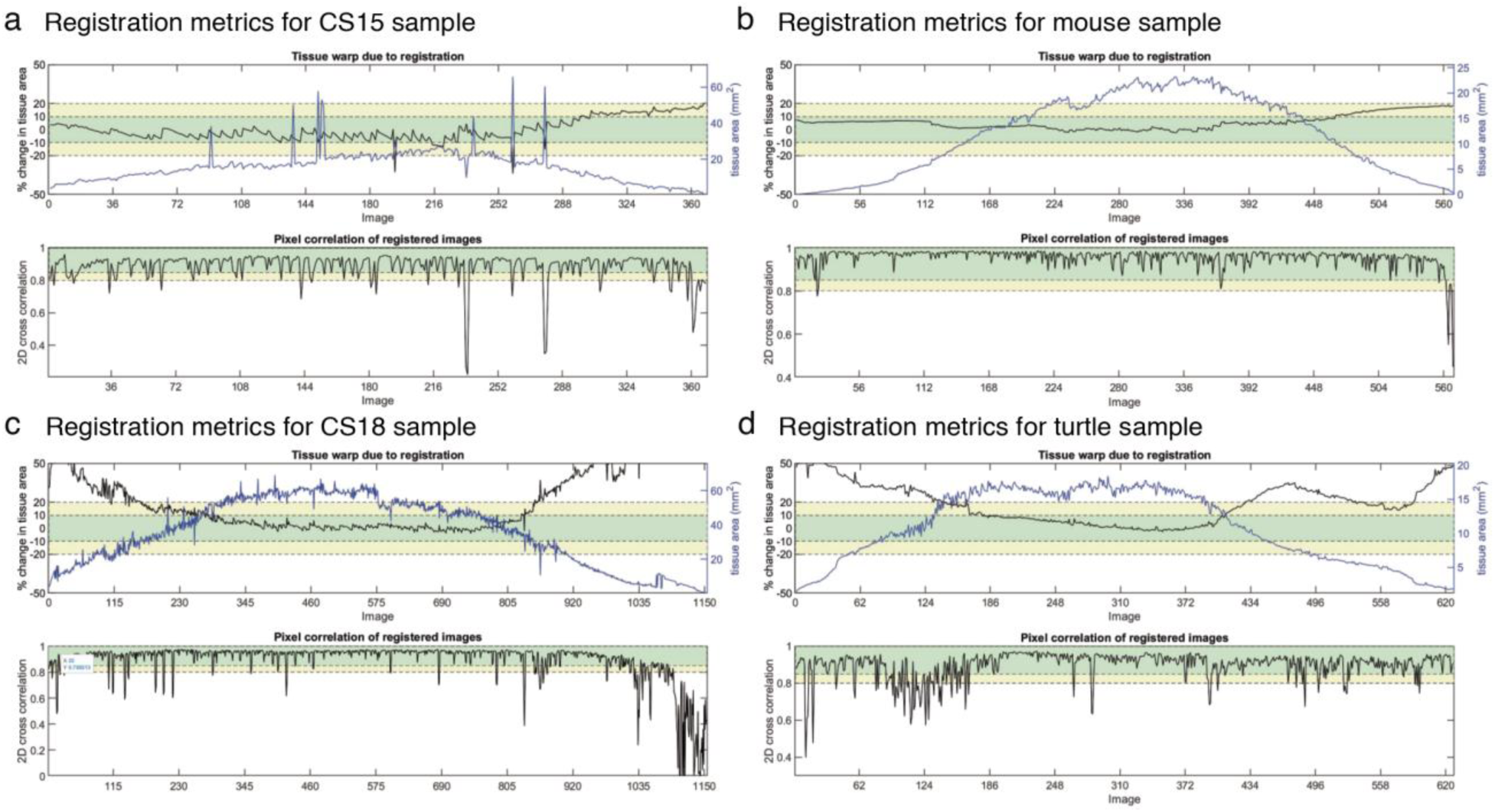
Quantitative assessment of 3D registration accuracy across embryonic specimens from multiple species. **(a)** Registration metrics for rhesus macaque CS15 sample demonstrate robust alignment performance across 380 serial sections. Upper panel shows tissue warp measurements with elastic registration. Lower panel displays pixel correlation coefficients between registered adjacent sections, maintaining values above 0.8 throughout the specimen with occasional drops corresponding to anatomical transitions or sectioning artifacts. **(b)** Registration metrics for mouse embryo sample spanning 565 sections show similar alignment stability. **(c)** Registration metrics for CS18 sample across 1,150 sections demonstrate scalability to larger datasets. Despite increased specimen complexity, warp corrections remain bounded within acceptable parameters. **(d)** Registration metrics for turtle embryo sample covering 620 sections confirm cross-species applicability of the alignment pipeline. Tissue warp measurements remain stable throughout registration.

**Fig. S3.**
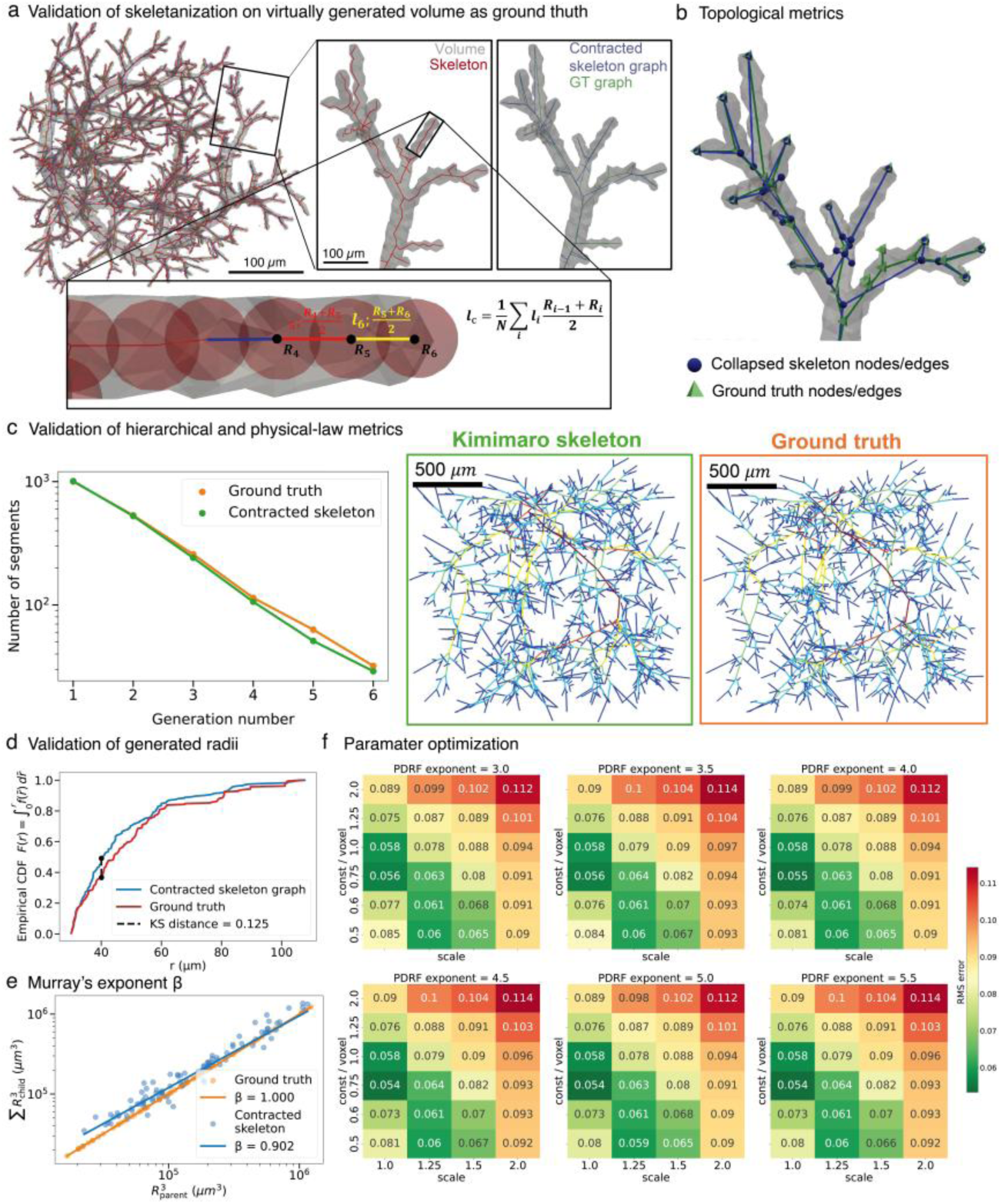
Validation of 3D skeletonization algorithms and topological analysis methods. **(a)** Validation of skeletonization on virtually generated volume from ground truth graph skeleton. Upper panel shows synthetic vascular network with known topology used for algorithm benchmarking. Detailed comparison between TEASAR-derived skeleton (red) and contracted skeleton graph ground truth (gray) demonstrates high fidelity preservation of branching topology and centerline accuracy. Lower panel illustrates radius estimation methodology using overlapping sphere fitting, where local radius (R_i_) is computed as the average of adjacent sphere radii weighted by segment length. **(b)** Topological metrics comparison between collapsed skeleton nodes/edges (blue circles) and ground truth nodes/edges (green triangles) shows excellent correspondence across network hierarchies. Quantitative analysis confirms >95% topological preservation with minimal spurious branch generation. **(c)** Validation of hierarchical and physical-law metrics using Kimimaro skeletonization algorithm. Left panel demonstrates exponential scaling relationships for both ground truth (orange) and contracted skeleton (green) with nearly identical scaling exponents across generation numbers. Right panels show direct visual comparison between Kimimaro-derived skeleton and ground truth networks, confirming preservation of essential branching patterns and connectivity. **(d)** Validation of generated radii through cumulative distribution analysis. Kolmogorov-Smirnov test (p=0.125) confirms no significant difference between contracted skeleton graph and ground truth radius distributions, validating the sphere-fitting radius estimation approach. **(e)** Murray’s exponent ß validation demonstrates consistent scaling relationships between parent and daughter vessel radii. Ground truth yields ß=1.000±0.05, while contracted skeleton produces ß=0.902±0.08, confirming preservation of fundamental scaling laws governing vascular optimization. **(f)** Parameter optimization heatmaps for probability density function exponents ranging from 3.0 to 5.5 across multiple scale factors. Color-coded error matrices identify optimal parameter combinations (dark green regions) that minimize reconstruction error while preserving topological accuracy. Systematic parameter sweeps ensure robust performance across diverse network morphologies and species-specific variations.

**Fig. S4.**
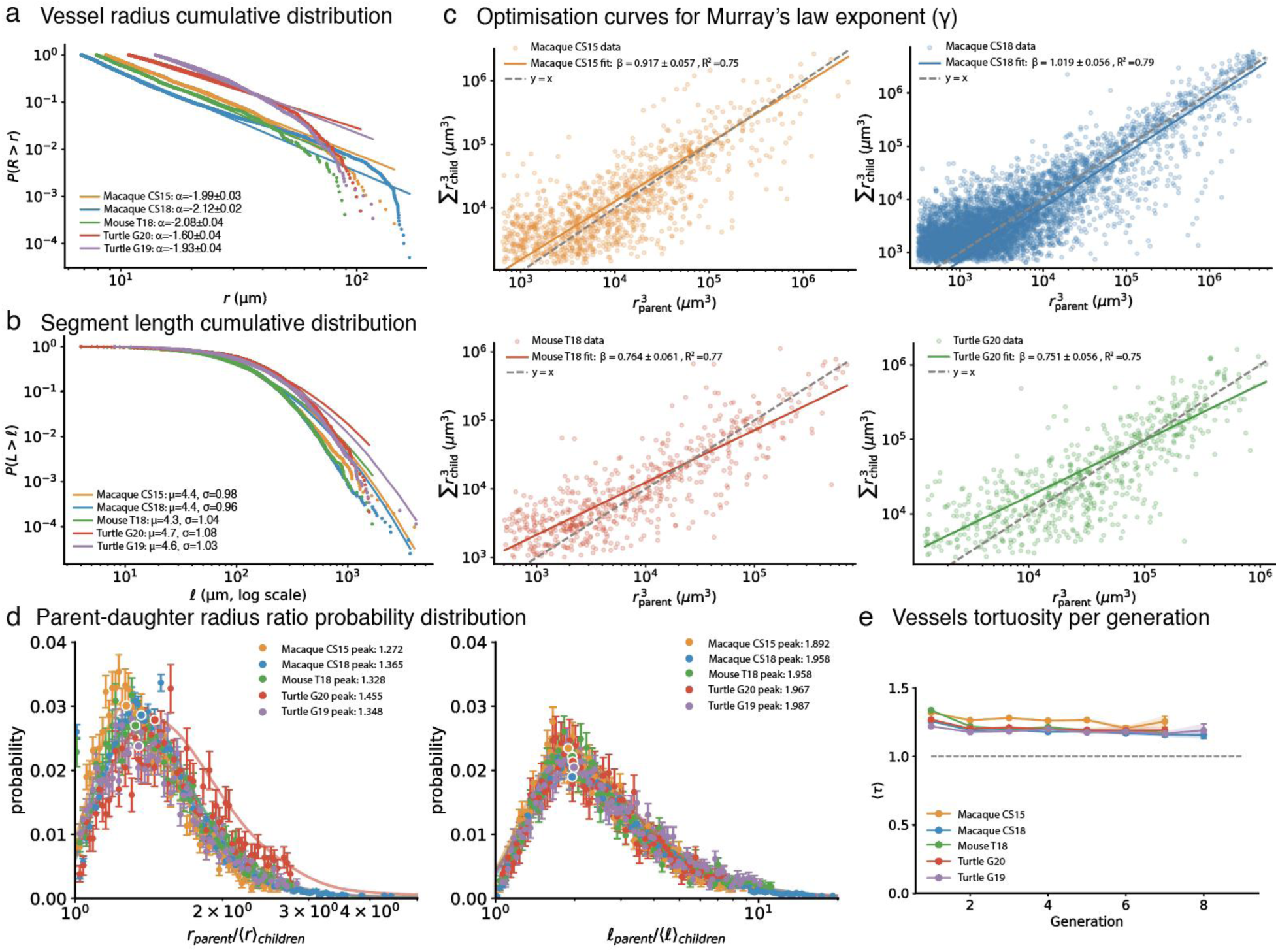
Morphometric analysis reveals universal scaling relationships in vascular network organization across species and developmental stages. **(a)** Vessel radius cumulative probability distributions P(R > r) with corresponding power-law scaling exponents: CS15 (α = -1.99 ± 0.03), and CS18 (α = -2.12 ± 0.02), T18 (α = -2.08 ± 0.04), turtle G19 (α = -1.60 ± 0.04). **(b)** Segment length cumulative probability distributions P(L > l) on log-log axes, revealing log-normal scaling with species-specific parameters: Macaque CS15 (μ = 4.4, σ = 0.98), and CS18 (μ = 4.4, σ = 0.96), mouse T18 (μ = 4.3, σ = 1.04), turtle G19 (μ = 4.7, σ = 1.08). **(c)** Murray’s law validation through parent-daughter radius cubed relationships for CS15 (β = 0.917 ± 0.057, R² = 0.75), CS18 (β = 1.019 ± 0.056, R² = 0.79), T18 (β = 0.764 ± 0.061, R² = 0.77), and G19 (β = 0.751 ± 0.056, R² = 0.75). Dashed lines indicate theoretical Murray’s law prediction (y = x). **(d)** Left: parent-daughter radius ratio probability distributions peak. Right: parent-daughter length ratio probability distributions demonstrate consistent branching geometry across species. **(e)** Vessel tortuosity remains stable across generations (≈1.2–1.3) for all specimens, indicating consistent vessel path efficiency throughout the vascular hierarchy. Error bars represent standard deviations.

**Fig. S5.**
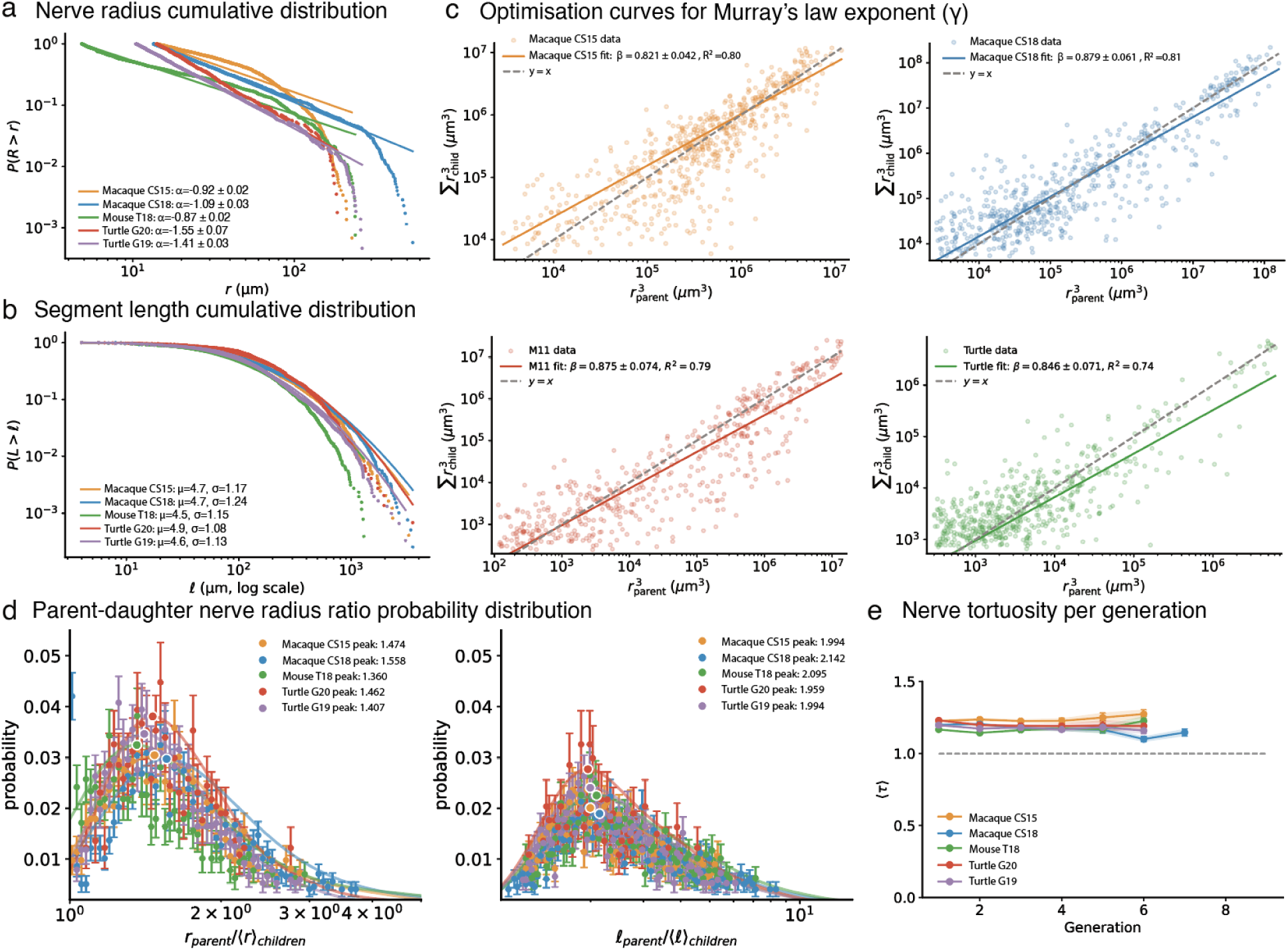
Morphometric analysis of nerve network reveals universal scaling relationships organization across species and developmental stages. **(a)** Nerve radius cumulative probability distributions P(R > r) on log-log axes, revealing power-law scaling with species-specific exponents: Macaque CS15 (α = -0.92 ± 0.02), CS18 (α = -1.09 ± 0.03), mouse T18 (α = -1.11 ± 0.04), turtle G20 (α = -1.55 ± 0.07), and G19 (α = -1.41 ± 0.03). **(b)** Nerve segment length cumulative probability distributions P(L > l) with corresponding log-normal scaling parameters: CS15 (μ = 4.7, σ = 1.17), CS18 (μ = 4.7, σ = 1.24), T18 (μ = 5.2, σ = 1.25), G20 (μ = 4.9, σ = 1.08), and G19 (μ = 4.6, σ = 1.13). **(c)** Murray’s law validation through parent-daughter radius cubed relationships for CS15 (β = 0.821 ± 0.042, R² = 0.80), CS18 (β = 0.879 ± 0.061, R² = 0.81), T18 (β = 0.875 ± 0.074, R² = 0.79), and turtle (β = 0.845 ± 0.071, R² = 0.74 and β = 0.879 ± 0.061, R² = 0.81). **(d)** Left: parent-daughter nerve radius ratio probability distributions peak near optimal branching ratios, with observed peaks: CS15 (1.474), CS18 (1.558), T18 (1.960), G20 (1.662), and G19 (1.407). Right: parent-daughter length ratio probability distributions demonstrate consistent branching geometry across species, with peaks: CS15 (1.994), CS18 (2.142), T18 (2.095), G20 (1.959), and G19 (1.994). **(e)** Nerve tortuosity remains stable across generations (≈1.0–1.2) for all specimens, indicating consistent nerve path efficiency throughout the nerve hierarchy. Error bars represent standard deviations.

## SUPPLEMENTAL TABLES

**Table S1.** Morphometric parameters of vascular and nervous system networks across different species.

| Sample | Nervous System |  |  |  | Vascular Network |  |  |  |
| --- | --- | --- | --- | --- | --- | --- | --- | --- |
| | Segments | $L_{min}$<br>($\mu m$) | $L_{max}$<br>($\mu m$) | $L_{chord}$<br>($m$) | Segments | $L_{min}$<br>($\mu m$) | $L_{max}$<br>($\mu m$) | $L_{chord}$<br>($m$) |
| Macaque CS15 | 2465 | 4.0 | 3256.4 | 0.5 | 10274 | 4.0 | 3962.5 | 1.2 |
| Macaque CS18 | 3968 | 4.0 | 3537.5 | 0.9 | 39859 | 4.0 | 3636.3 | 4.7 |
| Mouse T18 | 2576 | 4.0 | 1299.1 | 0.4 | 8786 | 4.0 | 1661.3 | 0.8 |
| Turtle G20 | 1476 | 4.0 | 3537.3 | 0.3 | 4705 | 4.0 | 1554.3 | 0.8 |
| Turtle G19 | 3054 | 4.0 | 3050.2 | 0.5 | 8779 | 8.0 | 4101.1 | 1.4 |

**Table S2.** Power-law fitting parameters and ranges for vascular network distributions.

| Sample | $\alpha$ | $x_{min}$ | $n_{tail}$ |
| --- | --- | --- | --- |
| Macaque CS15 | $2.99 \pm 0.03$ | 8.65 | 3781 |
| Macaque CS18 | $3.12 \pm 0.02$ | 6.80 | 19847 |
| Mouse T18 | $3.08 \pm 0.04$ | 7.89 | 2422 |
| Turtle G20 | $2.60 \pm 0.04$ | 10.80 | 2045 |
| Turtle G19 | $2.93 \pm 0.04$ | 14.00 | 2926 |

**Table S3.** Power-law fitting parameters and ranges for nervous system network distributions.

| Sample | $\alpha$ | $x_{min}$ | $n_{tail}$ |
| --- | --- | --- | --- |
| Macaque CS15 | $1.92 \pm 0.02$ | 14.00 | 1470 |
| Macaque CS18 | $2.09 \pm 0.03$ | 13.44 | 1744 |
| Mouse T18 | $1.87 \pm 0.02$ | 4.83 | 1315 |
| Turtle G20 | $2.55 \pm 0.07$ | 14.00 | 565 |
| Turtle G19 | $2.41 \pm 0.03$ | 10.47 | 2167 |

**Table S4.** Horton ratios and fractal dimensions of vascular networks across different species.

| Sample | $R_N \pm \sigma$ | $R_r \pm \sigma$ | $D_r \pm \sigma$ |
| --- | --- | --- | --- |
| Macaque CS15 | $2.35 \pm 0.16$ | $1.37 \pm 0.05$ | $2.72 \pm 0.37$ |
| Macaque CS18 | $2.40 \pm 0.13$ | $1.23 \pm 0.03$ | $4.28 \pm 0.59$ |
| Mouse T18 | $2.25 \pm 0.08$ | $1.38 \pm 0.05$ | $2.49 \pm 0.30$ |
| Turtle G20 | $2.31 \pm 0.26$ | $1.32 \pm 0.03$ | $2.99 \pm 0.46$ |
| Turtle G19 | $2.21 \pm 0.04$ | $1.25 \pm 0.02$ | $3.62 \pm 0.30$ |

**Table S5.** Horton ratios and fractal dimensions of nervous system networks across different species.

| Sample | $R_N \pm \sigma$ | $R_r \pm \sigma$ | $D_r \pm \sigma$ |
| --- | --- | --- | --- |
| Macaque CS15 | $2.15 \pm 0.17$ | $1.53 \pm 0.04$ | $1.81 \pm 0.22$ |
| Macaque CS18 | $2.28 \pm 0.14$ | $1.79 \pm 0.07$ | $1.42 \pm 0.14$ |
| Mouse T18 | $2.42 \pm 0.21$ | $1.93 \pm 0.05$ | $1.35 \pm 0.14$ |
| Turtle G20 | $2.20 \pm 0.18$ | $1.50 \pm 0.04$ | $1.94 \pm 0.24$ |
| Turtle G19 | $2.43 \pm 0.17$ | $1.65 \pm 0.05$ | $1.78 \pm 0.18$ |

## METHODS

### Tissue acquisition and processing of embryonic samples

The rhesus macaque embryos specimens were obtained from the Oregon Health and Sciences University (OHSU) and Oregon National Primate Research Center (ONPRC). The Turtle specimens were obtained from Concordia Turtle Farm (Jonesville, LA). The Mouse embryos were obtained from Jackson laboratories. Specimens were processed into FFPE tissue blocks and exhaustively serial sectioned at 4 microns in sagittal plane and H&E stained (one every two sections)^37^. Unstained slides were stored in -20°C, under optimal humidity and vacuum conditions for prospective use. H&E-stained slides were scanned at 20x resolution (∼0.5 micron/pixel) using a Hamamatsu Nanozoomer S210 (Fig. 1a). NDPI files were converted to tiff images (1, 2, and 4 micron/pixel resolutions) using openslide^53^.

### CODA vascular and nervous system microanatomical WSI labelling of whole organisms

To label the vascular microanatomical components of entire embryos, we developed two CODA semantic segmentation models (Fig. 1a, Fig. S1a-f)^54,55^. One model labelled the endothelium walls of the blood vessels, smooth muscle, muscle, and condensed mesenchyme in all WSIs (Fig. S1a-b). The second model was designed to automatically annotate the lumen of blood vessels and categorize all other tissues as a single class called tissue (Fig. S1c-d). Models were combined to maximize the segmentation accuracy of vessels in the embryonic samples. InterpolAI was used to generate missing images to restore microanatomical connectivity^56^. Expert histological annotations were generated as ground truth masks and validated against semantically segmented images generated by our trained model. This validation process resulted in an overall training accuracy exceeding 90% (Fig. S1a,c). To label the nervous system and peripheral nerves of all specimens, the same procedure of the CODA pipeline was performed (Fig. 3).

### Alignment of 2D WSIs of entire embryos into 3D maps

Combination of global rigid and local elastic image registrations allowed reconstruction of microanatomical structures of specimens into 3D volume^54^. Alignment was applied to images subtyping the vascular and nervous systems. Registration accuracy was quantitatively assessed using tissue warp measurements and pixel correlation coefficients between adjacent sections (Fig. S2). Optimal registration maintained correlation values between adjacent sections above 0.8, and percentage change in tissue area should be ideally less than 20% of change (Fig. S2a-d). Registration tissue variations were imputed into calculations as error variations.

### Network skeletonization and topological analysis

Three-dimensional vascular and nerve networks were extracted from segmented volumes using the TEASAR skeletonization algorithm (Fig. 1c, Fig. 6)^38^. Centerline representations preserved essential branching topology while enabling quantitative morphometric analysis (Fig. S3a-b). Topological validation employed synthetic vascular networks generated using VascuSynth with known ground truth topology, enabling direct comparison between extracted skeletons and ground truth graphs (Fig. S3a-b). Local vessel radii were computed using overlapping sphere fitting methods, where radius estimates incorporated weighted averaging across adjacent segments (Fig. S3a). Topological validation employed synthetic vascular networks generated using VascuSynth^43^ with known ground truth topology, enabling direct comparison between extracted skeletons and ground truth graphs (Fig. S3a-b). Systematic optimization of TEASAR/Kimimaro algorithm^38^ parameters across scale and const/voxel settings minimized RMS error relative to ground truth across multiple PDRF exponents (Fig. S3f). Validation confirmed preservation of hierarchical branching relationships across generations (Fig. S3c), accurate radius distribution recovery (KS distance = 0.125; Fig. S3d), and faithful reproduction of Murray’s law scaling relationships (extracted β = 0.902 vs. ground truth β = 1.000; Fig. S3e).

### Power-law and log-normal distribution fitting

Power-law distributions were fitted to empirical vessel and nerve segment data using maximum likelihood estimation (MLE)^57^, implemented via the Python powerlaw package^58^. MLE on continuous, unbinned data was employed rather than linear regression on log-log binned histograms, as binning introduces bias through subjective bin width selection and overweights noisy fluctuations in the distribution tail. The complementary cumulative distribution function (CCDF) was used for visualization as it provides a statistically accurate representation of the unbinned MLE model fit without requiring binning or smoothing.

The lower bound (x_min_) from which the power-law scaling relationship holds was determined dynamically by sweeping through the data to identify the threshold minimizing the Kolmogorov-Smirnov (KS) distance between empirical and analytical distributions (Table S2, Table S3). For vessel segment radii, fitted parameters were: CS15 (α=2.99±0.03, x_min_=8.65, n_tail_=3781), CS18 (α=3.12±0.02, x_min_=6.80, n_tail_=19847), T18 (α=3.08±0.04, x_min_=7.89, n_tail_=2422), GB20 (α=2.60±0.04, x_min_=10.80, n_tail_=2045), and GB19 (α=2.93±0.04, x_min_=14.00, n_tail_=2926). For nerve segment radii: CS15 (α=1.92±0.02, x_min_=14.00, n_tail_=1470), CS18 (α=2.09±0.03, x_min_=13.44, n_tail_=1744), T18 (α=1.87±0.02, x_min_=4.83, n_tail_=1315), GB20 (α=2.55±0.07, x_min_=14.00, n_tail_=565), and GB19 (α=2.41±0.03, x_min_=10.47, n_tail_=2167).

For segment length distributions, log-normal models were fitted using unbinned MLE yielding scale (μ) and shape (σ) parameters. Unlike power-law fitting, the lower bound for log-normal distributions was fixed to the data minimum rather than dynamically optimized, as log-normal distributions characterize the bulk of the dataset rather than isolating the heavy tail.

### Hierarchical classification and generation analysis

Network hierarchies were classified using Strahler ordering systems adapted for vascular topology (Fig. 6c). Terminal segments received generation 1 classification, with generation numbers increasing toward central arterial trunks. Bifurcation ratios, segment length distributions, and diameter scaling relationships were computed for each hierarchical level (Fig. 2a-b, Fig. 4a-b, Fig. S4a). Generation-resolved statistics enabled cross-species comparison of fundamental scaling principles governing network organization (Fig. S4a).

### Murray’s law analysis and optimization assessment

Vascular network optimization was evaluated through Murray’s law analysis, which predicts optimal diameter relationships minimizing metabolic costs of fluid transport (Fig. 2d-e, Fig. 4d-e). Parent-daughter vessel diameter relationships were fitted to power-law scaling:

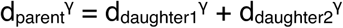

To quantify the uncertainty of γ*, we employed empirical bootstrapping. By repeatedly resampling our observed parent-child bifurcations with replacement, we constructed 1,000 simulated network datasets of the exact same size as the original. Because this process randomly duplicates some bifurcations while omitting others, it effectively simulates the natural biological variation expected across distinct biological specimens. We re-optimized the least-squares error curve for each of these 1,000 independent iterations to establish a robust 95% confidence interval for the exponent estimate. Optimal γ values were determined through least-squares fitting with theoretical predictions (γ=3.0 for fluid transport, γ=1.5-2.0 for electrical conduction). Nerve networks were analyzed using modified Murray exponents appropriate for signal transmission optimization rather than fluid transport (Fig. 4d-e). Validation using synthetic networks confirmed accurate parameter estimation across diverse topologies (Fig. S3d-e).

### Cross-species morphometric comparison and scaling analysis

Quantitative morphometric features were extracted across species including segment length distributions, radius scaling relationships, and topological complexity measures (Fig. 2, Fig. 3, Fig. 4). Power-law fitting was applied to identify universal scaling exponents using maximum likelihood estimation (Fig. 2c, Fig. 4c, Fig. S4). Kolmogorov-Smirnov tests evaluated goodness-of-fit for proposed scaling relationships. Cross-species conservation was assessed through comparative analysis of scaling parameters and hierarchical organization principles across human, mouse, and turtle embryonic specimens (Fig. 3, Fig. 6).

### Statistical considerations

Registration accuracy was quantified using normalized cross-correlation and mutual information metrics (Fig. S2). Morphological distributions (radius, length) were analyzed using Maximum Likelihood Estimation (MLE) to fit power-law and log-normal models. For power-law distributions, the optimal lower-bound threshold (x_min_) was determined by minimizing the Kolmogorov-Smirnov distance between the empirical data and the fitted model, with parameter uncertainties for the scaling exponents determined analytically. Network optimality was evaluated by determining the bifurcation exponent (γ) through minimization of the log-transformed least-squares error of the generalized Murray’s Law (r_parent_^γ^ = Σr_children_^γ^) using bounded scalar optimization routines. Model selection between power-law and log-normal distributions employed likelihood ratio tests with goodness-of-fit assessed through Kolmogorov-Smirnov statistics. Statistical significance was defined as p<0.05 with confidence intervals determined where appropriate.

## Code and data availability statement

The data analyzed here is available from the corresponding author upon request. The code used to generate the 3D tissue maps is available on the following GitHub: https://github.com/ashleylk/CODA. The code to skeletonize the vascular and nerve networks and to perform the analysis shown here is upon publication.

## Author contributions

A.F., A.L.K. and D.W. conceived the project. A.F., P.G., W.F., L.D., J.L., G.B., A.L.K, and D.W. collected and processed the specimens. A.F., M.C, C.O., V.Q., S.J., M.W., M.R.G, O.J.T.M, G.B, B.M., S.X.S conducted the analysis. A.F., A.L.K., and D.W. wrote the first draft of the manuscript, which all authors edited and approved.

## SUPPLEMENTAL MATERIAL

### Theoretical framework for vascular network optimization

Murray’s law for vascular networks emerges from minimization of the combined viscous pumping power and metabolic maintenance power for steady incompressible flow in passive pipe networks. The hydraulic resistance scales with vessel radius (r) and length (l), where the volumetric flux (Q) of a fluid with viscosity μ relates to the pressure drop Δp as 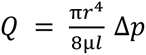. The mechanical power required to sustain flow is 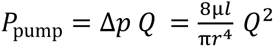. Since blood is biologically active, metabolic energy proportional to vessel volume maintains viability. Denoting the metabolic maintenance rate per unit volume as λ, the metabolic power is 𝑃𝑚𝑒𝑡 = λ π𝑟^2^𝑙. The total power required to flow biologically active blood through the network is therefore 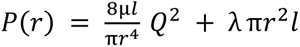. For biologically efficient networks, the flow rate Q minimizes P(r), such that 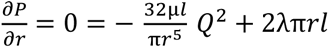. Solving for Q yields 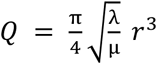, hence flow carried by a vessel scales with the cube of its radius. At bifurcations where the parent vessel divides into N daughter branches with radii, mass conservation together with the flow-radius relationship yields Murray’s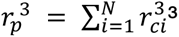.

Empirical validation of this cubic scaling relationship in our datasets (Fig. 7).

### Theoretical framework for nerve network optimization

For unmyelinated cylindrical axons (representative of our fetal samples) with radius r and length Δx, Kirchhoff’s law states 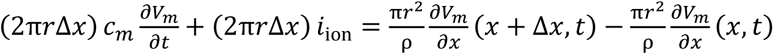, where c_m is specific membrane capacitance, ρ is intracellular resistivity, and i_ion is trans-membrane ionic current density. Dividing by 2πrΔx and taking the limit Δx → 0 yields the cable equation 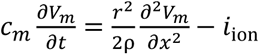.

The longitudinal current conserved at bifurcations is 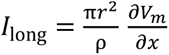. At branch points where a parent axon (radius r_1_) splits into two daughters (radii r_2_, r_3_), given that the travelling potential waveform is identical across each branch node and local resistivity remains unchanged around branching nodes, current conservation 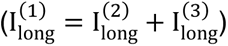 yields 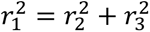. For symmetric bifurcations (𝑟_2_ = 𝑟_3_ ≡ 𝑟*_d_*), this reduces to 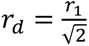, representing a Murray-type law with exponent 𝛾 =2. Repeated binary branching under this distribution predicts a size distribution 𝑁(𝑟) ∝ 𝑟^−2^ for axon populations: 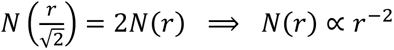.

Empirical measurements of bifurcation exponents and radius distributions consistent with these theoretical predictions (Fig. 8).

